# Comprehensive Comparative Genomics Reveals Over 50 Phyla of Free-living and Pathogenic Bacteria are Associated with Diverse Members of the Amoebozoa

**DOI:** 10.1101/2020.10.03.324772

**Authors:** Yonas I. Tekle, Janae M. Lyttle, Maya G. Blasingame

## Abstract

The association of bacteria with microbial eukaryotes has been extensively studied. Among these the supergroup Amoebozoa containing predominantly amoeboid unicellular protists has been shown to play an important ecological role in controlling environmental bacteria. Amoebozoans not only graze bacteria but also serve as a safe niche for bacterial replication and harbor endosymbiotic bacteria including dangerous human pathogens. Despite their importance, only a few lineages of Amoebozoa have been studied in this regard. Amoebozoa encompasses lineages of extreme diversity in ecology, morphology and evolutionary history. The limited amoebozoans studied are not representative of the high diversity known in the supergroup, and could undermine our understanding of their role as key players in environmental ecosystems and as emerging public health threats. In this research, we conducted a comprehensive genomic and transcriptomic study with expansive taxon sampling by including representatives from the three known clades of the Amoebozoa. We used culture independent whole culture and single cell genomics maintained in our laboratory cultures, and additionally published RNA-Seq data to investigate the association of bacteria with diverse amoebozoans. Relative to current published evidence, we recovered the largest number of bacterial phyla (57) and pathogen genera (49) associated with the Amoebozoa. Using single cell genomics we were able to determine up to 24 potential endobiotic bacterial phyla, some potentially endosymbionts. This includes the majority of multidrug-resistant pathogens designated as major public health threats. Our study demonstrates amoebozoans are associated with many more phylogenetically diverse bacterial phyla than previously recognized. It also shows that all amoebozoans are capable of harboring far more dangerous human pathogens than presently documented, making them of primal public health concern.

## Introduction

The study of microbial interactions is a complex and fascinating field of research ^1-3^. Microorganisms occupy diverse ecological niches and are usually found in large communities that result in inherent interactions. Coevolutionary processes have been shaping these interactions, which gave rise to various types of adaptation, specialization and establishment of temporary and stable (obligate) associations ^2,4-6^. Understanding microbial interactions have profound evolutionary implications; among other notable insights, it has contributed to our understanding of the origin of eukaryotic cells ^7^, ecosystem health and function ^8^ as well as disease and pathogen evolution ^9-11^. While the biodiversity of microbes is generally poorly understood, many examples of well-established associations are known among various microbes ^12^. Among these, the interactions of bacteria with protists (single-cell eukaryotes) have been a subject of immense scientific interest and substantial investigations ^9-11,13^. Protists comprise some of the most important primary grazers of environmental bacteria. They play an integral role in major biogeochemical and ecological processes of microbial food webs, substantially contributing to nutrient recycling and energy transfer to higher trophic levels both in aquatic and terrestrial ecosystems ^2,14^ Furthermore, many animal and human pathogenic bacteria are directly or indirectly associated with protists. Several studies have shown that many bacteria, including some that are well-known multidrug resistant (e.g. *Legionella*), are capable of evading digestion by protists ^3,15-17^ These bacteria use protist hosts as safe haven to reproduce and as intermediate agents to infect their final hosts. Many examples of this type of relationship are known in ciliates ^13^, flagellates ^18^ and amoeboids ^3,9,14^ In this study, we will focus on the association of bacteria with the predominantly amoeboid supergroup, Amoebozoa.

The association of bacteria with Amoebozoa has been mostly studied from two representatives, *Acanthamoeba* and *Dictyosletlium* ^19-23^. These two amoebozoans are extensively studied as models in many important cellular processes and pathogenesis ^10,20,24-27^. Some reports on association with bacteria are also available in a few other amoebozoan genera (e.g. *Vermamoeba, Platyamoeba/Vannella*) ^17,20,28-30^. These studies demonstrated that amoebozoans are both grazers and hosts of some bacterial epibionts (attached to the surface of the amoebozoan) and endobionts (within the cytoplasm of the amoebozoan), the latter including dangerous human pathogens. Amoebozoans have been implicated as training ground for emerging pathogens and vehicles for their dispersal ^4,20^. These studies also gave insights on mechanism of pathogen evasion and host defense ^16,20,26,31^. Despite these major advances in the field, the number of amoebozoans examined for association with bacteria remain limited; and the studied lineages are not representative of the extremely diverse groups currently recognized within the supergroup. Amoebozoa encompasses members characterized by diverse morphology, ecology, behavior and life cycle ^32-35^. The limited taxa used to study association with bacteria, undoubtedly has missed the vast diversity of bacteria that could potentially be associated with the Amoebozoa. Consequently, this under sampling hampers our knowledge of the positive contributions, and impact, that amoebozoans might have on the environment; and their role in major public health concerns.

Over ten bacterial pathogens (in humans and other eukaryotes) belonging to the commonly discovered five bacterial phyla (Proteobacteria, Bacteroidetes, Chlamydiae, Firmicutes and Actinobacteria) have been reported in the Amoebozoa ^4,9,17,20,24,28,29^. Additionally, some less known bacteria phyla (e.g. Dependentiae), and unclassified or novel bacterial lineages, have been reported to form temporary or stable endosymbiotic associations with some amoebozoans ^6,36^. These reports are mostly based on culture-dependent studies, which focus on the microbiome of bacteria that can be cultured concurrently with the target amoebozoan. Culture-dependent studies fail to capture those bacteria that are unculturable under conventional laboratory conditions and with established culture media. Studies that used a culture-independent approach also suffer from taxon sampling, or they are limited to specialized or specific environments ^29,37^. In order to capture the complete microbiome of the Amoebozoa-associated bacteria, we used a culture-independent, comprehensive genomic approach and surveyed 49 samples (38 species) covering most known lineages of Amoebozoa. The samples come from the three major clades of Amoebozoa, consisting of lineages of different morphology, ecology and behavior ^32^. We used large genetic sampling, including genome data derived from whole culture and single cells maintained in our laboratory and transcriptome data obtained from prior published research. We assessed the impact of sampling and culturing conditions on the types and number of bacteria discovered. Our study reveals 57 bacterial phyla, including 49 known human pathogenic genera, associated with the various members of the Amoebozoa. Our study reports the largest number of associated bacteria, including new phyla and pathogen genera, not reported in previous studies. Our findings reinforce previous reports that showed Amoebozoa as a major grazer of environmental bacteria, and host of many bacterial endosymbionts, some that pose a threat to public health. This study also lays foundation for further investigations on mechanisms of predator-prey relationships, evasion of host defense (immunity) and forms and types of associations of newly discovered epi- and endobionts, some that are symbiotic and others that are internalized pathogenic bacteria.

## Results

### Overall Composition of Amoebozoa Associated Bacteria

Taxonomic assignment of the various genetic datasets analyzed, combining genome data generated in this study with transcriptomes from previous studies, yielded a large number of amoebozoan-associated bacteria phyla with overall similar taxonomic compositions across the three clades of Amoebozoa. A total of 57 bacterial phyla were discovered from all of the datasets examined (Fig. 1, Tables S1-S3). Since the majority of bacterial phyla, 56, were found in the whole culture RNA-Seq dataset, we will focus our comparison among the clades of Amoebozoa based on this dataset mostly (Fig. 2). One additional phylum besides others was found in the whole culture genome dataset (Table S3). Discosea, with the highest number of taxa analyzed in whole culture RNA-seq dataset, contained 52 bacterial phyla, while Evosea and Tubulinea had 44 and 39 phyla, respectively (Fig. 1, Table S1). Among these discovered phyla, 33 phyla are shared among the three clades (Fig. 2A). While the bacterial taxon sampling for Tubulinea in the transcriptome data is smaller than Evosea and Discosea, the latter two clades shared more bacterial phyla between them (i.e. 9), when compared to the phyla that they each mutually shared with Tubulinea (i.e. 1 and 3, respectively) as shown in Fig. 2A. We also found some bacterial phyla specifically associated with each clade; namely, 7 in Discosea, 2 in Tubulinea and 1 in Evosea (Fig. 2A). However in future research, the specific bacterial phyla recovered in each clade might change with more taxon sampling, and in relation to the nature of the acquired data. For instance, two samples from the same species, *C. minus*, in the whole culture genome data showed variation in the number of bacterial phyla recovered and shared (Table S3). This indicates that a thorough and even sampling is required to make such comparisons. Overall, phyla recovered were proportional to data size and taxon sampling (Fig. 1, Tables S1-S3).

**Figure 1.**
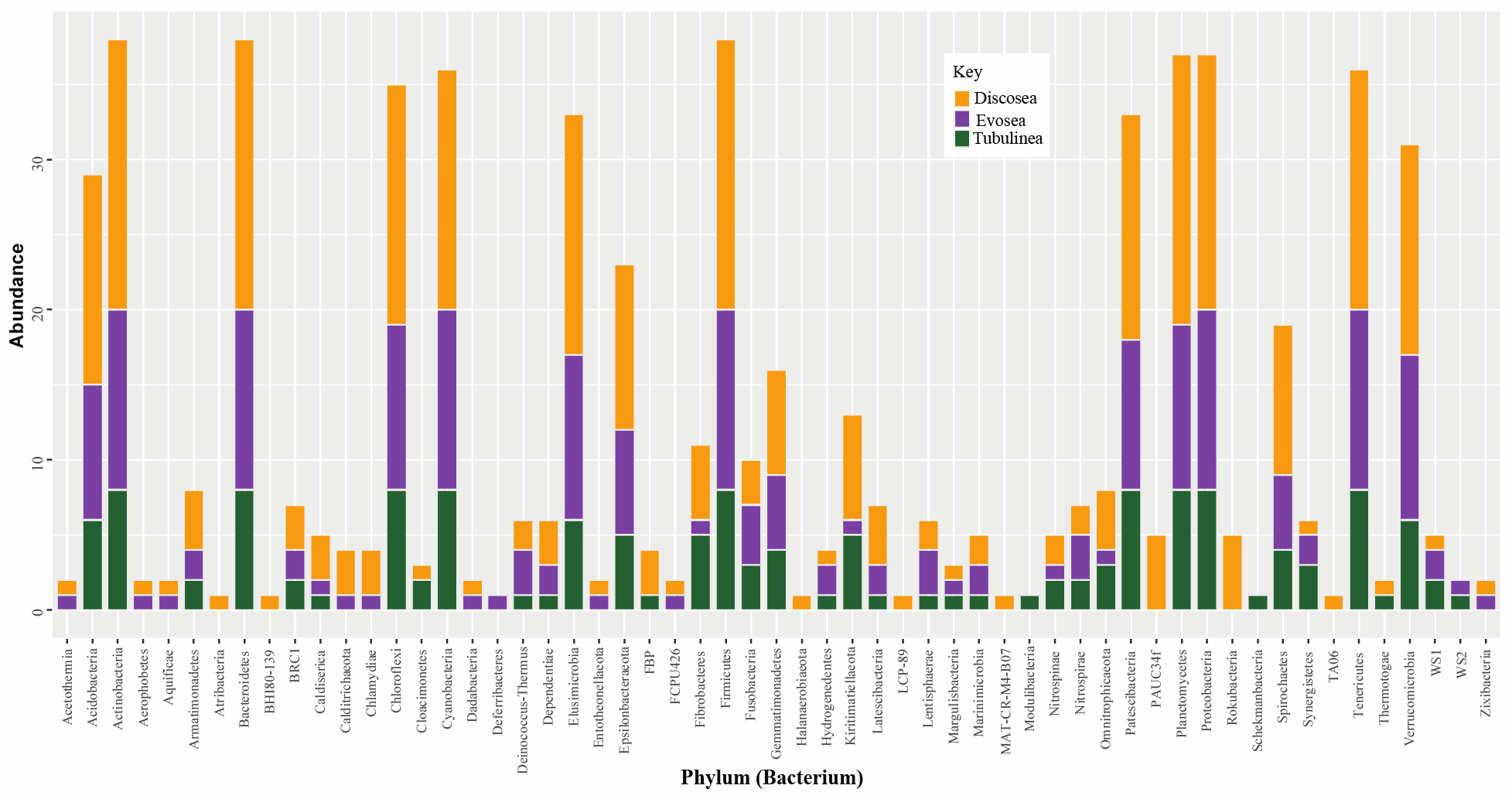
A distribution of genera representing 57 Bacterial phyla discovered in the three major clades of Amoebozoa across all datasets analyzed.

**Figure 2.**
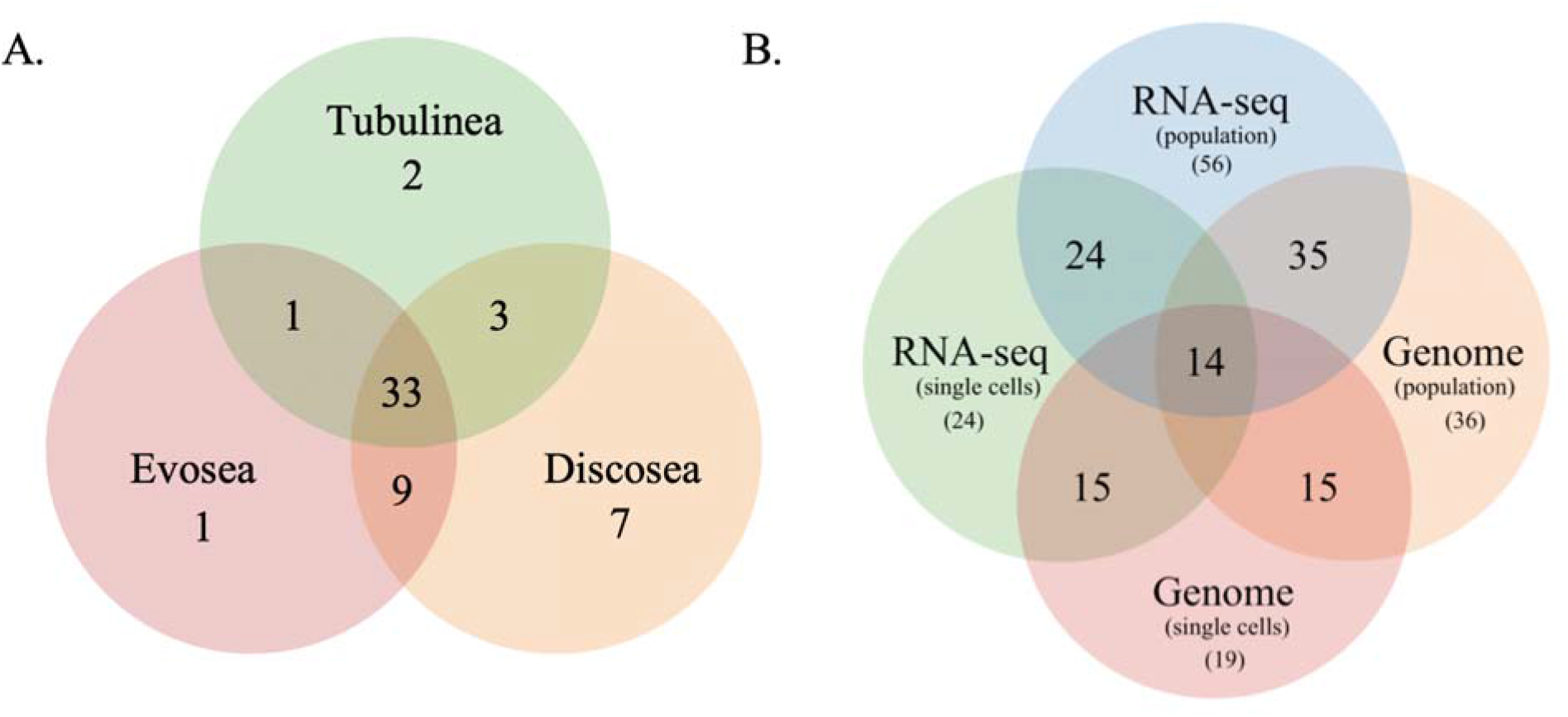
Venn diagram showing bacterial phyla shared among the three major clades of Amoebozoa of the whole culture RNA-Seq data (A) and among the four types of datasets analyzed (B).

The total number of genera and their representation differed by bacterial phyla in our datasets. The most abundant bacterial phylum recovered in all datasets and amoebozoan clades is Proteobacteria (Tables 1, Tables S1-S3). Class Gammaproteobacteria, a subdivsion of Proteobacteria, was represented by a higher number of genera and total number of sequences that were representative for its genera (Tables 1, S1-S5). Other bacterial phyla were represented by over 1000 sequences for the genera recovered, including Bacteroidetes and Firmicutes (Table 1). Generally, a higher number of sequences representing a given phylum were observed in the whole culture genome data (Table S3).

**Table 1.** List of potential endosymbiont (pathogens) bacterial phyla and their abundance (total number of sequences) found in all or at least 3 datasets analyzed.

| Phylum | Present | Total | Pathogen |
| --- | --- | --- | --- |
| Proteobacteria | 4/4 | 16501 | **Alpha** - *Ehrlichia, Rickettsia*; **Beta** - *Bordetella, Burkholderia, Neisseria*; **Epsilon** – *Campylobacter*; **Gamma** – *Acinetobacter, Coxiella, Enterobacter, Escherichia, Francisella, Haemophilus, Klebsiella, Legionella, Proteus, Pseudomonas, Salmonella, Serratia, Shigella, Vibrio, Yersinia* |
| Bacteroidetes | 4/4 | 2028 | *Chryseobacterium, Porphyromonas, Prevotella* |
| Firmicutes | 4/4 | 1773 | *Bacillus, Clostridium, Enterococcus, Faecalibacterium, Staphylococcus, Streptococcus* |
| Patescibacteria | 4/4 | 452 | - |
| Actinobacteria | 4/4 | 404 | *Actinomyces, Corynebacterium, Mycobacterium, Nocardia, Propionibacterium, Rhodococcus, Rothia, Trueperella* |
| Cyanobacteria | 4/4 | 378 | - |
| Chloroflexi | 4/4 | 347 | - |
| Tenericutes | 4/4 | 267 | *Mycoplasma, Ureaplasma* |
| Planctomycetes | 4/4 | 233 | - |
| Verrucomicrobia | 4/4 | 189 | - |
| Acidobacteria | 4/4 | 140 | - |
| Epsilonbacteraeota | 4/4 | 104 | - |
| Nitrospirae | 4/4 | 33 | - |
| Gemmatimonadetes | 4/4 | 32 | - |
| Elusimicrobia | 3/4 | 171 | - |
| Spirochaetes | 3/4 | 53 | *Brachyspira, Borrelia, Leptospira, Treponema* |
| Dependentiae | 3/4 | 36 | - |
| Armatimonadetes | 3/4 | 19 | - |
| Fibrobacteres | 3/4 | 18 | - |
| Omnitrophicaeota | 3/4 | 14 | - |
| Marinimicrobia/SA R406 | 3/4 | 13 | - |
| BRC1 | 3/4 | 13 | - |
| Synergistetes | 3/4 | 9 | *Acetomicrobium, Cloacibacillus, Synergistes* |
| Latescibacteria | 3/4 | 9 | - |

Two pathogens containing phyla, Chlamydiae (*Neochlamydia*) and Fusobacteria (*Fusobacterium*), have been detected in this study but were only found in one of our datasets.

### Comparison of Data types and Potential Endosymbiont Bacterial Phyla

The four data types analyzed yielded bacterial phyla that are commonly shared among samples and amoebozoan clades analyzed (Figs. 2, S1, Tables S1-S5). We observed some variations in taxonomic breadth and the total number genera recovered depending on data type and taxon sampling size (Figs. 2, S1, Tables S1-S5). As mentioned above all except one bacterial phylum reported here were present in whole culture RNA-Seq datasets (Table S1). While the large number of bacterial phyla in the whole culture RNA-Seq dataset can be partly attributed to the size of taxon sampling used for this dataset, these results clearly indicates that RNA-Seq is a good data source for this type of study. The whole culture genome data is represented by two independent samples from a single species, *C. minus* (Table S3). A total of 36 bacterial phyla were recovered from these samples, 35 of these are shared with the whole culture RNA-Seq dataset (Fig. 2B). The single cell genome data yielded the lowest number, 19 bacterial phyla (Table S3), after the single cell RNA-Seq data (24 phyla) (Tables S2). Using the four datasets we were able to identify 14 potential endobionts/epibionts by taking a subset of the bacterial phyla discovered in each dataset (Fig. 2B). Use of single cells datasets, both genome and RNA-Seq, primarily aimed at reducing bacteria contamination from external environment enabled us to deduce these 14 putative endobionts/epibionts. A total of 24 potential endobionts/epibionts phyla can be recognized if we considered taxa shared among three datasets i.e. all the phyla discovered in single cell RNA-Seq dataset (Fig. 2B, Table 1). Among these seven putative endobionts phyla (5 shared in all and 2 shared among 3 datasets, Table 1) included members (human pathogen genera) previously reported to associate with or found in the cytoplasm of amoebozoan hosts ^3,9,22,24,31,46^.

In order to assess the impact of culturing techniques and types of bacteria that may be associated due to difference in the environment of isolation and types of food sources used between labs, we compared RNA-Seq data of three taxa sequenced in two different labs. Our comparison showed similar total number of bacterial phyla recovery but with some differences in the number of overlapping phyla (Table S1). The variation of non-overlapping phyla in these three pairs of species ranged from 5-7. This observed difference using the RNA-Seq data is smaller compared to the variation observed in the number of non-overlapping phyla found in the genome data samples (Table S3). The whole culture genome data used two samples from the same species that were cultured under the same conditions. These two samples had 9 non-overlapping bacterial phyla, which indicate that other technical factors, such as sample quality and sequencing, might affect the recovery rate of overlapping bacterial community in samples of the same species.

### Human pathogenic Bacterial phyla and genera associated with Amoebozoa

Our survey of literature for bacterial human pathogens yielded over 60 genera spanning 10 phyla (Table S4). We used this list to investigate the presence of pathogenic genera in our datasets (Tables S4-S5). Of the 67 bacterial human pathogenic genera surveyed, 49 pathogens were found belonging to 9 different phyla (Figs. 3, S1, Table 1, Tables S4, S5). The number of pathogens recovered in the three clades, Discosea (39 pathogens), Evosea (35 pathogens) and Tubulinea (33 pathogens), were similar despite taxon sampling differences in the whole culture RNA-Seq dataset (Fig. 3, Table S4). We also recovered a similar set of pathogens among the four datasets (whole culture and single cell genome and transcriptome datasets, 30-44 pathogens); except, the single cell genome dataset had a lower (11) number of pathogens (Fig. S1, Tables S4, S5). These eleven pathogens discovered in our single cell genome dataset belonged to bacterial phyla that were shown to be putative endosymbionts (see above, Fig. 2B, Tables 1, S5).

**Figure 3.**
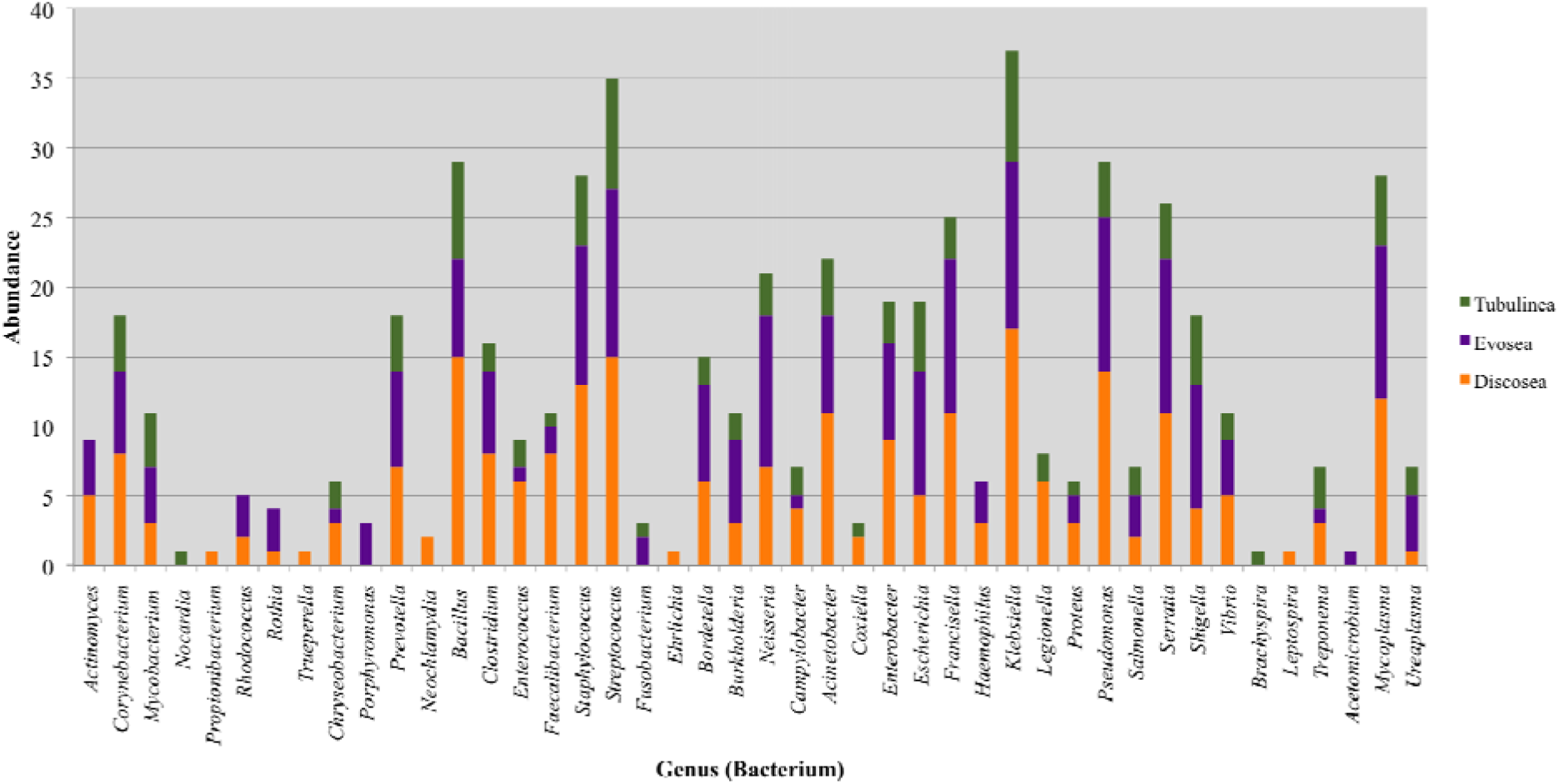
Distribution of the 44 pathogenic bacterial genera discovered in the three major clades of Amoebozoa in the whole culture RNA-Seq data.

The top three phyla with the highest number of pathogenic bacterial genera recovered include Proteobacteria, Actinobacteria and Firmicutes (Table 1). Among the classes of Proteobacteria, Gammaproteobacteria had the largest number (15 pathogen genera) compared to any group analyzed (Table 1). Of the nine pathogen containing phyla five were found in at least 3 datasets, while two including Chlamydiae (*Neochlamydia*) and Fusobacteria (*Fusobacterium*), were rare and only recovered in one dataset (Table 1, Fig. S1).

## Discussion

### Large Amoebozoa Associated Bacterial Phyla Recovered

Our study using whole culture and single cell genomics and transcriptomics recovered the largest number of bacterial phyla that are potentially associated with the supergroup Amoebozoa to date. The majority of the bacterial phyla recovered in our analysis of the amoebozoans are newly reported here for the first time (Fig. 1, Tables S1-S3). We also found well known and common amoebozoan-associated bacterial phyla reported in previous studies ^3,4,6,9,15,24,28-30,37^ The large and taxonomically diverse discovery of amoebozoans associated bacterial phyla in this study could be attributed to the comprehensive taxon sampling and molecular genetic approach employed. We analyzed amoebozoans characterized by diverse ecology, behavior and evolutionary history that represented the three major clades of the Amoebozoa. We used monoclonal cultures of amoebozoans isolated directly from nature or acquired from culture collection agents ^32,33,35^. Research methods using monoclonal cultures typically include addition of food bacteria (e.g., *E. coli* or *Klebsiella*); but once the culture starts to advance, it is common to see more bacterial communities, besides food bacteria, growing among the amoebozoan cells. Amoebozoans are known to carry undigested food bacteria vertically through generations. These food bacteria are used presumably as seeds to be conserved for potential replenishment within new environments encountered by the amoebozoan, and then harvested as food; this behavior led some to metaphorically call amoebozoans, ‘farmers’ ^20,47-49^. Therefore, the bacteria found in monoclonal samples analyzed likely reflect a bacterial community that might be expected to occur naturally in nature; although we cannot rule out that some are acquired from contamination during laboratory culture as for example from contact with instruments used in culturing or from air-borne bacteria introduced from the laboratory environment. The taxonomic composition of bacteria found in amoebozoans grown in different labs, or obtained from different culture collection agents, in the RNA-Seq data were similar (Table S1). The consistent recovery of similar bacterial phyla across different amoebozoan samples and taxonomic groups, that we have found in our analyses for this research study, also indicates that all bacterial lineages discovered in our analysis are potentially associated with the Amoebozoa, and may mitigate against possible contamination from sources largely derived from the laboratory environment. While the confirmation and type of association of the newly discovered bacteria awaits further investigation, our study reinforces amoebozoans as key players in controlling environmental bacteria through grazing. Our study also shows that Amoebozoa harbor more taxonomically diverse bacteria, with 64% of the 89 bacterial phyla in SILVA database recovered, than previously reported.

The large taxonomic sampling of amoebozoans in our study was made possible by the use of transcriptome data. In recent phylogenomic studies, a large number of RNA-Seq datasets have been generated in the Amoebozoa ^32,33,35^. These transcriptome data are generated using a standard approach that selects polyadenylated RNA (polyA) in RNA samples, which selects against bacterial contaminant transcripts that are typically poorly polyadenylated ^50,51^. However, transcriptome data collected from amoebozoans using this approach typically contains large bacterial transcripts and some ribosomal genes ^32,33,35^. While contamination by bacteria in transcriptome data has been reported in axenic culture, or in species that do not normally feed or associate with bacteria (likely contamination from environment) ^52^, the close association of bacteria (food and endosymbiont) with amoebozoans exacerbates the potential for contamination of transcriptomes even more. We took advantage of this, and used the 16S bacterial ribosomal genes found in amoebozoan RNA-Seq data to assess bacterial association with the Amoebozoa. Despite the potential limitation that transcriptome data might have for our study, the aggregate number of bacterial phyla recovered from transcriptome sequencing was comparable in taxonomic coverage to the whole culture genome data (Fig. 2). As expected, the number of genomic representations of the discovered phyla in the whole culture genome data was higher than the transcriptome data (Table S1-S3), which indicates that transcriptome data might to an extent underrepresent the actual abundance of associated bacterial populations. Our results support the utility of transcriptome data to study association of bacteria with amoebozoans or other similar protists. Though a conservative estimate, transcriptome data has some advantages over genome data due to lower cost and ease in acquiring it. Moreover, transcriptome data can provide additional information on the nature of an association by providing physiological data (profile of expressed genes) among interacting species ^53^.

In addition to the rich sources of transcriptome data as discussed above, the use of whole culture and singe cell genomics, as used in our laboratory culture studies reported here, enabled us to assess potential bacterial endobionts (possibly including epibionts) associated with the Amoebozoa. Using this approach we identified 14-24 potential endobionts/epibionts bacterial phyla (Fig. 2B, Table 1). Our list includes bacteria phyla whose members were previously shown to form true endosymbiotic relationship in some amoebozoans ^6,9,28,54,55^. However, a more thorough approach including single cell genome and cytological data, such as use of fluorescently labeled oligonucleotide probes (e.g., Horn et al., 2000), is needed to establish true endosymbiotic relationships with Amoebozoa. Nonetheless, the recovery of known endosymbiotic bacteria in our analysis gives credence to the reliability of our approach to identify potential endosymbiotic bacteria candidates that can be studied further. It should be noted that some amoebozoans are selective bacterial predators ^56-58^. The combination of single cell genomics and transcriptomics approaches used here is a promising method of analyzing selective feeding on bacteria by protists; e.g., a recent study demonstrated the utility of transcriptome data for selective feeding in a ciliate lineage ^53^.

### Pathogenic bacteria associated with the Amoebozoa

The association of pathogenic bacteria with some members of Amoebozoa has been investigate in great detail ^3,4,20,21,26,59^. Most of the association of pathogenic bacteria described with amoebozoans is facultative, but some permanent associations are also known ^6,28,46^. While most associations are transient and harmless, some bacterial infections (e.g. *Legionella*), leading to lysis of amoebozoan cells, have been reported ^4,60^. In facultative associations, the pathogenic bacteria can use the amoeba cell as a safe niche to reproduce, or intermediate host, or even as a vehicle for dispersal or population reservoir ^4,21^. Some recent studies have proposed that amoebozoans could serve as an ‘interim training ground’ to develop intracellular survival strategies before becoming a human pathogen due to the similarity in mechanism of phagocytosis (phagolysosome) within mammalian macrophages ^4,16,27^ Most of the known pathogenic bacteria associated with Amoebozoa so far come from the studies that used only a few amoebozoan species, which are not necessarily reflective of pathogens that can be harbored by various groups in the supergroup of Amoebozoa. In this study, we discovered 49 pathogenic bacterial genera belonging to 9 phyla, the highest report so far (Table 1). The number and distribution of pathogenic genera across the three major groups of Amoebozoa were comparable despite differences in taxon sampling among them (Figs. 3, S1). Our list includes previously reported common pathogen bacterial phyla ^20,59^ in addition to the large number of pathogens newly discovered in this study (Tables 1, S4). Congruent with previous studies, the most dominant pathogen-containing phylum is Proteobacteria. One of its subdivisions, class Gammaproteobacteria, comprised more than 50% of the pathogenic genera identified in this study (Table 1). Interestingly one of the bacterial pathogen phylum, Chlamydiae, frequently recovered in previous studies ^28,46,61^ was very rare and only found in one of our data sets. Several of the pathogenic bacteria found associated with amoebozoans are studied from anthropogenic habitats (e.g. cooling towers, hospitals, humidifier aerosols, drinking water, spas or swimming pools) ^23,29,30,37,54^. The representation of some pathogen-containing phyla might be affected by habitat examined. Nevertheless, our results demonstrate that all amoebae are potential carriers of bacterial pathogens both in nature or anthropogenic environments. All of the multidrug resistant genera (except *Helicobacter*) found in this study are listed and categorized by CDC and WHO as urgent, and various levels of threats and concerns. Among these are *Acinetobacter, Clostridium, Enterococci, Neisseria, Campylobacter, Pseudomonas, Salmonella, Mycoplasma, Streptococcus, Bordetella* that were found in the amoebozoans we examined (see Table 1). This makes some Amoebozoa that are associated with potential or acknowledged human pathogens a major public health threat.

## Materials and Methods

### Whole Culture and Single Cell Genomics

We used various approaches to investigate bacteria associated with amoebozoans. Association of bacteria with their host can be internal endobionts (some endosymbionts) or external those that are epibionts attached to the surface of the cell and those that are freely present in cultures that are potentially available to be engulfed as a food source. In order to capture all associated bacteria in diverse monoclonal cultures of amoebozoans in our laboratory, we used molecular data collected using two approaches. The first set of genetic data collected consisted of community genomic DNA extracted from actively growing cultures of amoebozoans; and from the bacterial community typically found in monoclonal or newly isolated species maintained in our laboratory cultures. The second genetic data is derived from single amoebozoan cells, individually picked out of our laboratory cultures. The main difference between these two approaches is that the first approach, *whole culture*, is aimed at collecting large quantities of DNA from a monoclonal population without little consideration to bacteria contamination from the culture; while the second approach, *single cell*, is aimed at minimizing bacterial contamination from the surrounding environment.

In the single cell approach, amoebozoan cells including *Cochliopodium minus, Stratorugosa tubuloviscum, Trichosphaerium* sp. and *Amoeba proteus* were individually picked using mouth pipetting techniques and transferred to a clean glass slide to wash off bacteria (other microbial eukaryotes (food or prey) in *A. proteus* culture) to reduce contamination of freely growing bacteria (other contaminants) from the culture. This step does not necessarily remove epibionts that are tightly bound to the cell surface but it greatly minimizes free (loosely bound) bacteria growing in culture. *Stratorugosa tubuloviscum* and *C. minus* were grown in plastic Petri dishes with bottled natural spring fresh water (Deer Park®, Nestlé Corp. Glendale, CA, USA) with added autoclaved grains of rice as an organic nutrient source to support bacterial growth as prey for the amoebozoans. The marine *Trichosphaerium* sp. was grown under a similar condition as above in artificial seawater. *Amoeba proteus* was purchased from Ward’s Science culture collection (wardsci.com) and was cultured with mixed bacteria and other microbial eukaryote food sources. Cleaned individual cells (5-10) were transferred into 0.2-mL PCR tubes and genome amplified using REPLI-g Advanced DNA Single Cell Kit (Qiagen Hilden, Germany). For the whole culture approach, genomic DNA was extracted from a large number of *Cochliopodium minus* (syn. *C. pentatrifurcatum* ^38,39^ cells in culture dishes (50 Petri dish cultures) using MagAttract high-molecular-weight (HMW) DNA kit (Qiagen, MD), following the manufacturer’s instructions. This method includes gentle cell lysis, releasing high molecular weight DNA and its efficient isolation and purification by concentration on DNA-binding, surface coated magnetic beads. Genome sequencing was performed using 10X genomics (for whole culture DNA) and Oxford Nanopore (ONP) (for both single cells and whole culture DNA) following the manufacturers’ protocol. Genome data from 10X genomics and ONP were assembled using Supernova v2.1.1 ^40^ and Minimap2-Miniasm-Racon genome assembly pipeline ^41-43^, respectively. For ONP genome data we used Filtlong version 0.2.0 (https://github.com/rrwick/Filtlong) to filter reads with length shorter than 200 and quality score less than 5. Porechop version 0.2.4 (https://github.com/rrwick/Porechop) was used to remove ONP sequencing adapters added during the sequencing.

### Whole Culture and Single Cell Transcriptome Data

Based on preliminary analysis that showed amoebozoan transcriptomes contained large bacterial transcripts and some ribosomal genes, we analyzed RNA-Seq from previous publications collected in a similar manner as above ^32,33,35,44^. The whole culture RNA-Seq dataset included a total of 35 species (15 discoseans, 12 evoseans, and 8 tubulinids) with three additional duplicate samples from Discosea sequenced in two different labs ^32,33,35^. These discosean duplicate samples were included in the analysis to examine the effects of culturing methods and environment on the number and composition bacterial community recovered. The single cell RNA-Seq dataset was represented by 5 samples obtained from *Cochliopodium minus* ^44^. Data collection, sequencing and assembly of transcriptome data of these diverse amoebozoans, representing the three main clades of Amoebozoa (Discosea, Evosea, and Tubulinea) of the whole culture and single cell RNA-Seq datasets, are described in Kang et al. ^32^ and Tekle et al. ^33,35^, and Tekle et al. ^44^, respectively. Some good quality transcriptomes whose origin was not certain or is collected using a combination of single cell and whole culture are placed in the whole culture RNA-Seq dataset. All transcriptomes used for single cell RNA-Seq dataset are collected in our laboratory under similar experimental conditions ^44^.

### Taxonomic Assignment of Amoebozoa Associated Bacterial Sequence Data

Taxonomic assignment of the assembled contigs (>300 pbs) from genome and transcriptome data was performed with Kraken 2 ^45^. This program’s sequence algorithm classifies sequences by mapping k-mer to the lowest common ancestor (LCA) of all the datasets containing the given k-mer in the specified database. The 16S database, SILVA, was chosen for this analysis and taxonomic classification was done to a genus level. Kraken 2 was run with default settings locally in an interactive session on XSEDE server, a supercomputing platform (http://xsede.org). To obtain broad evidence of amoebozoan-associated bacteria, we analyzed a total of 49 samples (genome and transcriptome data) of amoebozoans, representing 38 species belonging to the three major clades of Amoebozoa. Similarly, we compared taxonomic composition results of genome and RNA-Seq data obtained using the whole culture and single approaches. Resulting data were further analyzed using R and Excel.

## Acknowledgments

This work is supported by the National Science Foundation EiR (1831958) and National Institutes of Health (1R15GM116103-02) to YIT. We would like to thank Fang Wang for technical assistance during data collection and analysis. O. Roger Anderson is thanked for his invaluable comments and edits on the manuscript.

## Supplementary Materials Captions

**Figure S1**. Distribution of the pathogenic bacterial genera discovered in the four datasets analyzed.

**Table S1**. Tally of bacterial genera in whole culture RNA-Seq dataset. All amoebozoans representing the three major clades including species pairs sequenced in different labs (shown in red font) are included.

**Table S2**. Tally of bacterial genera derived from single cells RNA-Seq dataset. For this analysis different samples from *Cochliopodium minus* were examined.

**Table S3**. Tally of bacterial genera derived from whole culture and single cells genome datasets.

**Table S4**. Tally of potential human pathogenic bacterial genera using the whole culture RNA-Seq data in amoebozoans representing the three major clades.

**Table S5**. Tally of potential human pathogenic bacterial genera in three datasets including Single cells and whole culture genome datasets and single cell RNA-Seq data.

## Reference

1 Gast, R. J., Sanders, R. W. & Caron, D. A. Ecological strategies of protists and their symbiotic relationships with prokaryotic microbes. Trends Microbiol 17, 563–569, doi:10.1016/j.tim.2009.09.001 (2009).

2 Braga, R. M., Dourado, M. N. & Araujo, W. L. Microbial interactions: ecology in a molecular perspective. Braz J Microbiol 47 Suppl 1, 86–98, doi:10.1016/j.bjm.2016.10.005 (2016).

3 Barker, J. & Brown, M. R. Trojan horses of the microbial world: protozoa and the survival of bacterial pathogens in the environment. Microbiology 140 (Pt 6), 1253–1259, doi:10.1099/00221287-140-6-1253 (1994).

4 Molmeret, M., Horn, M., Wagner, M., Santic, M. & Abu Kwaik, Y. Amoebae as training grounds for intracellular bacterial pathogens. Appl Environ Microbiol 71, 20–28, doi:10.1128/AEM.71.1.20-28.2005 (2005).

5 Hibbing, M. E., Fuqua, C., Parsek, M. R. & Peterson, S. B. Bacterial competition: surviving and thriving in the microbial jungle. Nat Rev Microbiol 8, 15–25, doi:10.1038/nrmicro2259 (2010).

6 Horn, M. et al. Obligate bacterial endosymbionts of Acanthamoeba spp. related to the beta-Proteobacteria: proposal of ‘Candidatus Procabacter acanthamoebae’ gen. nov., sp. nov. Int J Syst Evol Microbiol 52, 599–605, doi:10.1099/00207713-52-2-599 (2002).

7 Margulis, L. in Symbiosis in cell evolution: microbial communities in the Archean and Proterozoic eons 1–18 (W.H. Freeman, 1993).

8 Handley, K. M. Determining Microbial Roles in Ecosystem Function: Redefining Microbial Food Webs and Transcending Kingdom Barriers. mSystems 4, doi:10.1128/mSystems.00153-19 (2019).

9 Horn, M. & Wagner, M. Bacterial endosymbionts of free-living amoebae. The Journal of eukaryotic microbiology 51, 509–514, doi:10.1111/j.1550-7408.2004.tb00278.x (2004).

10 Erken, M., Lutz, C. & McDougald, D. The rise of pathogens: predation as a factor driving the evolution of human pathogens in the environment. Microb Ecol 65, 860–868, doi:10.1007/s00248-013-0189-0 (2013).

11 Amaro, F., Wang, W., Gilbert, J. A., Anderson, O. R. & Shuman, H. A. Diverse protist grazers select for virulence-related traits in Legionella. ISME J 9, 1607–1618, doi:10.1038/ismej.2014.248 (2015).

12 Mishustin, E. N. Microbial associations of soil types. Microb Ecol 2, 97–118, doi:10.1007/BF02010433 (1975).

13 Gong, J. et al. Protist-Bacteria Associations: Gammaproteobacteria and Alphaproteobacteria Are Prevalent as Digestion-Resistant Bacteria in Ciliated Protozoa. Front Microbiol 7, 498, doi:10.3389/fmicb.2016.00498 (2016).

14 Anderson, O. R. The role of Bacterial-based Protist Communities in Aquatic and Soil Ecosystems and the Carbon Biogeochemical Cycle, with Emphasis on Naked Amoebae. Acta Protozoologica 51, 209–221. (2012).

15 Schmitz-Esser, S. et al. The genome of the amoeba symbiont “Candidatus Amoebophilus asiaticus” reveals common mechanisms for host cell interaction among amoeba-associated bacteria. Journal of bacteriology 192, 1045–1057, doi:10.1128/JB.01379-09 (2010).

16 Best, A. M. & Abu Kwaik, Y. Evasion of phagotrophic predation by protist hosts and innate immunity of metazoan hosts by Legionella pneumophila. Cell Microbiol 21, e12971, doi:10.1111/cmi.12971 (2019).

17 Greub, G. & Raoult, D. Microorganisms resistant to free-living amoebae. Clin Microbiol Rev 17, 413–433, doi:10.1128/cmr.17.2.413-433.2004 (2004).

18 Foster, R. A., Carpenter, E. J. & Bergman, B. Unicellular cyanobionts in open ocean dinoflagellates, radiolarians, and tintinnids: ultrastructural characterization and immuno-localization of phycoerythrin and nitrogenase. Journal of Phycology 42, 453–463 (2006).

19 Clarke, M. Recent insights into host-pathogen interactions from Dictyostelium. Cell Microbiol 12, 283–291, doi:10.1111/j.1462-5822.2009.01413.x (2010).

20 Thewes, S., Soldati, T. & Eichinger, L. Editorial: Amoebae as Host Models to Study the Interaction With Pathogens. Front Cell Infect Microbiol 9, 47, doi:10.3389/fcimb.2019.00047 (2019).

21 Strassmann, J. E. & Shu, L. Ancient bacteria-amoeba relationships and pathogenic animal bacteria. PLoS Biol 15, e2002460, doi:10.1371/journal.pbio.2002460 (2017).

22 Benavides-Montano, J. A. & Vadyvaloo, V. Yersinia pestis Resists Predation by Acanthamoeba castellanii and Exhibits Prolonged Intracellular Survival. Appl Environ Microbiol 83, doi:10.1128/AEM.00593-17 (2017).

23 Garcia, M. T., Jones, S., Pelaz, C., Millar, R. D. & Abu Kwaik, Y. Acanthamoeba polyphaga resuscitates viable non-culturable Legionella pneumophila after disinfection. Environ Microbiol 9, 1267–1277, doi:10.1111/j.1462-2920.2007.01245.x (2007).

24 Alibaud, L. et al. Pseudomonas aeruginosa virulence genes identified in a Dictyostelium host model. Cell Microbiol 10, 729–740, doi:10.1111/j.1462-5822.2007.01080.x (2008).

25 Dallaire-Dufresne, S., Paquet, V. E. & Charette, S. J. [Dictyostelium discoideum: a model for the study of bacterial virulence]. Can J Microbiol 57, 699–707, doi:10.1139/w11-072 (2011).

26 Bozzaro, S. & Eichinger, L. The professional phagocyte Dictyostelium discoideum as a model host for bacterial pathogens. Curr Drug Targets 12, 942–954, doi:10.2174/138945011795677782 (2011).

27 Cirillo, J. D. et al. Intracellular growth in Acanthamoeba castellanii affects monocyte entry mechanisms and enhances virulence of Legionella pneumophila. Infect Immun 67, 4427–4434 (1999).

28 Horn, M. et al. Neochlamydia hartmannellae gen. nov., sp. nov. (Parachlamydiaceae), an endoparasite of the amoeba Hartmannella vermiformis. Microbiology 146 (Pt 5), 1231–1239, doi:10.1099/00221287-146-5-1231 (2000).

29 Gomez-Alvarez, V., Revetta, R. P. & Santo Domingo, J. W. Metagenomic analyses of drinking water receiving different disinfection treatments. Appl Environ Microbiol 78, 6095–6102, doi:10.1128/AEM.01018-12 (2012).

30 Fields, B. S. et al. Characterization of an axenic strain of Hartmannella vermiformis obtained from an investigation of nosocomial legionellosis. J Protozool 37, 581–583, doi:10.1111/j.1550-7408.1990.tb01269.x (1990).

31 Segal, G. & Shuman, H. A. Legionella pneumophila utilizes the same genes to multiply within Acanthamoeba castellanii and human macrophages. Infect Immun 67, 2117–2124 (1999).

32 Kang, S. et al. Between a Pod and a Hard Test: The Deep Evolution of Amoebae. Molecular biology and evolution 34, 2258–2270, doi:10.1093/molbev/msx162 (2017).

33 Tekle, Y. I. et al. Phylogenomics of ‘Discosea’: A new molecular phylogenetic perspective on Amoebozoa with flat body forms. Molecular phylogenetics and evolution 99, 144–154, doi:10.1016/j.ympev.2016.03.029 (2016).

34 Tekle, Y. I. & Williams, J. R. Cytoskeletal architecture and its evolutionary significance in amoeboid eukaryotes and their mode of locomotion. R Soc Open Sci 3, 160283, doi:10.1098/rsos.160283 (2016).

35 Tekle, Y. I. & Wood, F. C. Longamoebia is not monophyletic: Phylogenomic and cytoskeleton analyses provide novel and well-resolved relationships of amoebozoan subclades. Molecular phylogenetics and evolution 114, 249–260, doi:10.1016/j.ympev.2017.06.019 (2017).

36 Deeg, C. M. et al. Chromulinavorax destructans, a pathogen of microzooplankton that provides a window into the enigmatic candidate phylum Dependentiae. PLoS pathogens 15, e1007801, doi:10.1371/journal.ppat.1007801 (2019).

37 Delafont, V., Brouke, A., Bouchon, D., Moulin, L. & Hechard, Y. Microbiome of free-living amoebae isolated from drinking water. Water Res 47, 6958–6965, doi:10.1016/j.watres.2013.07.047 (2013).

38 Tekle, Y. I., Anderson, O. R., Lecky, A. F. & Kelly, S. D. A New Freshwater Amoeba: Cochliopodium pentatrifurcatum n. sp. (Amoebozoa, Amorphea). The Journal of eukaryotic microbiology 60, 342–349, doi:10.1111/jeu.12038 (2013).

39 Tekle, Y. I. & Wood, F. C. A practical implementation of large transcriptomic data analysis to resolve cryptic species diversity problems in microbial eukaryotes. BMC evolutionary biology 18, 170, doi:10.1186/s12862-018-1283-1 (2018).

40 Weisenfeld, N. I., Kumar, V., Shah, P., Church, D. M. & Jaffe, D. B. Direct determination of diploid genome sequences. Genome research 27, 757–767, doi:10.1101/gr.214874.116 (2017).

41 Vaser, R., Sovic, I., Nagarajan, N. & Sikic, M. Fast and accurate de novo genome assembly from long uncorrected reads. Genome research 27, 737–746, doi:10.1101/gr.214270.116 (2017).

42 Li, H. Minimap2: pairwise alignment for nucleotide sequences. Bioinformatics 34, 3094–3100, doi:10.1093/bioinformatics/bty191 (2018).

43 Li, H. Minimap and miniasm: fast mapping and de novo assembly for noisy long sequences. Bioinformatics 32, 2103–2110, doi:10.1093/bioinformatics/btw152 (2016).

44 Tekle, Y. I., Wang, F., Heidari, A. & Stewart, A. J. Differential Gene Expression Analysis and Cytological Evidence Reveal a Sexual Stage of an Amoeba with Multiparental Cellular and Nuclear Fusion. bioRxiv, doi: doi: https://doi.org/10.1101/2020.06.23.166678 (2020).

45 Wood, D. E., Lu, J. & Langmead, B. Improved metagenomic analysis with Kraken 2. Genome biology 20, 257, doi:10.1186/s13059-019-1891-0 (2019).

46 Amann, R. et al. Obligate intracellular bacterial parasites of acanthamoebae related to Chlamydia spp. Appl Environ Microbiol 63, 115–121 (1997).

47 DiSalvo, S. et al. Burkholderia bacteria infectiously induce the proto-farming symbiosis of Dictyostelium amoebae and food bacteria. Proc Natl Acad Sci U S A 112, E5029–5037, doi:10.1073/pnas.1511878112 (2015).

48 Brock, D. A., Read, S., Bozhchenko, A., Queller, D. C. & Strassmann, J. E. Social amoeba farmers carry defensive symbionts to protect and privatize their crops. Nat Commun 4, 2385, doi:10.1038/ncomms3385 (2013).

49 Brock, D. A., Douglas, T. E., Queller, D. C. & Strassmann, J. E. Primitive agriculture in a social amoeba. Nature 469, 393–396, doi:10.1038/nature09668 (2011).

50 Nakazato, H., Venkatesan, S. & Edmonds, M. Polyadenylic acid sequences in E. coli messenger RNA. Nature 256, 144–146, doi:10.1038/256144a0 (1975).

51 Ohta, N., Sanders, M. & Newton, A. Poly(adenylic acid) sequences in the RNA of Caulobacter crescenus. Proc Natl Acad Sci U S A 72, 2343–2346, doi:10.1073/pnas.72.6.2343 (1975).

52 Strong, M. J. et al. Microbial contamination in next generation sequencing: implications for sequence-based analysis of clinical samples. PLoS pathogens 10, e1004437, doi:10.1371/journal.ppat.1004437 (2014).

53 Zou, S., Zhang, Q. & Gong, J. Comparative Transcriptomics Reveals Distinct Gene Expressions of a Model Ciliated Protozoa Feeding on Bacteria-Free Medium, Digestible, and Digestion-Resistant Bacteria. Microorganisms 8, doi:10.3390/microorganisms8040559 (2020).

54 Fritsche, T. R., Gautom, R. K., Seyedirashti, S., Bergeron, D. L. & Lindquist, T. D. Occurrence of bacterial endosymbionts in Acanthamoeba spp. isolated from corneal and environmental specimens and contact lenses. J Clin Microbiol 31, 1122–1126 (1993).

55 Proca-Ciobanu, M., Lupascu, G. H., Petrovici, A. & Ionescu, M. D. Electron microscopic study of a pathogenic Acanthamoeba castellani strain: the presence of bacterial endosymbionts. Int J Parasitol 5, 49–56, doi:10.1016/0020-7519(75)90097-1 (1975).

56 Singh, B. N. Selectivity in bacterial food by soil amoebae in pure mixed cultures and in sterilized soil. Ann. Appl. Biol. 28, 52–64. (1941).

57 Singh, B. N. The selection of bacterial food by soil amoebae, and the toxic effects of bacterial pigments and other products on soil protozoa. Brit. J. Exp. Path. 26, 316–325. (1945).

58 Ronn, R., McCaig, A. E., Griffiths, B. S. & Prosser, J. I. Impact of protozoan grazing on bacterial community structure in soil microcosms. Appl Environ Microbiol 68, 6094–6105, doi:10.1128/aem.68.12.6094-6105.2002 (2002).

59 Skriwan, C. et al. Various bacterial pathogens and symbionts infect the amoeba Dictyostelium discoideum. Int J Med Microbiol 291, 615–624, doi:10.1078/1438-4221-00177 (2002).

60 Molmeret, M., Bitar, D. M., Han, L. & Kwaik, Y. A. Disruption of the phagosomal membrane and egress of Legionella pneumophila into the cytoplasm during the last stages of intracellular infection of macrophages and Acanthamoeba polyphaga. Infect Immun 72, 4040–4051, doi:10.1128/IAI.72.7.4040-4051.2004 (2004).

61 Fritsche, T. R. et al. Phylogenetic diversity among geographically dispersed Chlamydiales endosymbionts recovered from clinical and environmental isolates of Acanthamoeba spp. Appl Environ Microbiol 66, 2613–2619, doi:10.1128/aem.66.6.2613-2619.2000 (2000).

